# EXPLOITING NEXT GENERATION SEQUENCING TO SOLVE THE HAPLOTYPING PUZZLE IN POLYPLOIDS: A SIMULATION STUDY

**DOI:** 10.1101/088112

**Authors:** Ehsan Motazedi, Richard Finkers, Chris Maliepaard, Dick de Ridder

**Affiliations:** Bioinformatics Group, Wageningen UR, The Netherlands; Wageningen UR Plant Breeding, The Netherlands

## Abstract

Haplotypes are the units of inheritance in an organism, and many genetic analyses depend on their precise determination. Methods for haplotyping single individuals use the phasing information available in Next Generation sequencing reads, by matching overlapping sNPs while penalizing post hoc nucleotide corrections made. Haplotyping diploids is relatively easy, but the complexity of the problem increases drastically for polyploid genomes, which are found in both model organisms and in economically relevant plant and animal species. While a number of tools are available for haplotyping polyploids, the effects of the genomic makeup and the sequencing strategy followed on the accuracy of these methods have hitherto not been thoroughly evaluated.

We developed the simulation pipeline *haplosim* to evaluate the performance of haplotype estimation algorithms for polyploids: HapCompass, HapTree and SDhaP, in settings varying in sequencing approach, ploidy levels and genomic diversity, using tetraploid potato as the model. Our results show that sequencing depth is the major determinant of haplotype estimation quality, that 1*kb* PacBio CCS reads and Illumina reads with large insert-sizes are competitive, and that all methods fail to produce good haplotypes when ploidy levels increase. Comparing the three methods, HapTree produces the most accurate estimates, but also consumes the most resources. There is clearly room for improvement in polyploid haplotyping algorithms.

## 1. Introduction

The advent of sequencing technology has had tremendous impact in genomics and genetics over the last years. The genome of *Arabidopsis thaliana*, the first completed plant genome published in 2000 [1], and the human (*Homo sapiens*) genome, published in 2003 [2], formed the basis for large-scale efforts to catalogue sequence variation found between individual genomes, in particular single nucleotide polymorphisms (SNPs) [3–5]. Subsequently, the 1001 *A*. *thaliana* genomes project [6] and the international HapMap project [7] sequenced a large number of individuals to discover haplotype blocks, i.e. genomic regions containing co-segregating SNPs. These haplotype blocks and their so-called haplotype tagging SNP markers, heterozygous SNPs whose alleles predict the presence of a certain haplotype, formed the basis for the development of high-density SNP arrays [8, 9], capable of determining rare or less-frequent genotypes in a population, which were used in a large number of studies relating SNPs to phenotypes such as ecological traits, diseases and disorders [10, 11].

Following the success of the *A*. *thaliana* and human genome projects, many animal and plant species were sequenced and genotyped, notably fruit fly (*Drosophila*) [12], chicken (*Gallus gallus*) [13], pig (*Sus scrofa*) [14], potato (*Solanum tuberosum*) [15], tomato (*S. lycopersicum*) [16] and hot pepper (*Capsium annuum*) [17] (the last three being plant genera within the *Solanacea* family). This has impacted not only fundamental genetics research, but has also revolutionized the fields of animal and plant breeding, by relating thousands of previously unknown genomic variants to physiological, morphological and economically important traits such as yield per generation and disease resistance [18, 19].

The introduction of high-throughput, relatively cheap and reliable next-generation sequencing (NGS) technologies made it possible to determine most of the variants directly within a single genome rather than using a pre-defined set of marker variants as proxies for the other variants [20]. Efficient tools have been developed to call variants in such sequencing data, e.g. FreeBayes [21] and GATK [22], and also to link variants on the same homologous chromosome, so-called haplotype phasing. However, while phasing of nearby variants occurring within the average NGS read length is relatively straightforward, long-range haplotyping using NGS data remains a challenge. Nevertheless haplotyping is important in many areas: in fundamental biology, to improve our understanding of genome structure, recombination and evolution [23]; in medicine, to obtain a full picture of the genetic variation in a population potentially linked to diseases and traits [24]; and in animal and plant breeding, to move from phenotype-based to genotype-based crossing and selection of individuals [25, 26].

In the diploid case, haplotyping algorithms aim to divide the reads into two complementary sets, each covering a specific region of, say, *n* heterozygous sites, so that the nucleotides are the same at the overlapping sites of the reads within each set, but different between the sets. The algorithmic challenge then is to take the occurrence of sequencing and variant calling errors into account [27–29]. Minimum Error Correction (MEC), the most prevalent approach, uses single base-flips for reads that conflict with both of the read sets, presumably due to sequencing or variant calling errors, to assign them to one of these. The aim is to find a configuration that requires a minimal number of such base flips [28,30]. This strategy is the basis of several diploid haplotyping algorithms [31–38].

For polyploid genomes, the problem could be formulated as dividing the reads into *k* > 2 groups, but the generalization from the diploid case is not straightforward. Unlike the diploid case, the knowledge of one haplotype does not automatically determine the phasing of the others. Besides, some haplotypes may be (locally) identical and thus several configurations could have the same MEC score. Moreover, the computational complexity of haplotype reconstruction increases rapidly with an increase in ploidy [39, 40]. Still, haplotype assembly for polyploids is highly relevant, as many interesting organisms have polyploid genomes and haplotyping will help unravel the range of the complex recombinations allowed by such genomes. Within the animal kingdom, triploidy and tetraploidy are observed in treefrog (*Xenopus laevis*) [41] and zebrafish (*Danio rerio*) [42], both important model organisms in evolutionary biology. Moreover, many economically important crops and ornamentals are polyploid, including tetraploid alfalfa (*Medicago sativa*), triploid banana (*Musa acuminata* × *M. balbisiana*), tetraploid leek (*Allium ampeloprasum*), tetraploid potato (*S. tuberosum*), tetraploid hard wheat (*Triticum durum*), hexaploid bread wheat (*T. aestivum*), tetraploid, hexaploid and octoploid strawberry species including *Faragaria moupinesis* (*k*=4), *F. moschata* (*k*=6), *F.* × *ananassa* (*k*=8) and several hybrid cotton (*Gossypium*, tetraploid or hexaploid) and rose (*Rosa*, tetraploid) species.

Here we review three state-of-the-art haplotyping algorithms applicable to polyploids: HapCompass [39, 43], HapTree [44] and SDhaP [40], and evaluate their accuracy through extensive simulations of random genomes and NGS reads. Using the highly heterozygous tetraploid potato (*S. tuberosum*) as a model, we generated random genomes using a realistic stochastic model with parameters SNP density and distribution of SNP dosages, i.e. the number of alternative alleles at each SNP site, derived from a recent genomic study of potato [45]. In addition, we simulated genomes at higher levels of ploidy with the same SNP density, as well as tetraploid genomes with different SNP densities and haplotype dosages, in order to investigate the effects of genome characteristics on the estimation. Moreover, we considered various sequencing depths, paired-end insert-sizes and sequencing technologies to quantify the impact of these parameters on the haplotyping. We provide guidelines to apply the haplotyping methods in practice, and show the characteristics of each method in various situations. The simulation pipeline is available as software package *haplosim*, which allows simulation for various sequencing approaches, genomic characteristics and variation models.

## 2. Material and Methods

We developed a multi-stage pipeline, *haplosim*, to simulate polyploid individuals with desired genomic characteristics, detect SNPs and their dosages for each individual, estimate the haplotypes and finally compare them to the simulated haplotypes using quantitative measures.

In the first step (Figure 1-A), random polyploid genomes are produced from a reference DNA sequence by introducing heterozygous regions containing bi-allelic SNPs, using the command-line tool *haplogenerator* that we developed for this purpose. Next, NGS reads are simulated for each produced individual using ART [46] and PBSIM [47] for Illumina and PacBio, respectively, and the reads are mapped back to their reference genome using *bwa-mem* [48]) (with the settings recommended in its manual for Illumina and PacBio reads). The alignments are pre-processed to generate BAM files and remove duplicates by samtools [49] and Picardtools [50], after which SNPs are called using FreeBayes [21] (Figure 1-B). The processed alignments, the reference and the VCF files are used in the haplotyping step by HapCompass [39, 43], HapTree [44] and SDhaP [40] to estimate the haplotypes using the phasing information from reads with at least two heterozygous SNPs (Figure 1-C). In the last step, the obtained estimates are compared to the original haplotypes by command-line tool *hapcompare* that we developed using several measures of estimation quality (Figure 1-D). These steps are explained in detail below.

**Figure 1.**
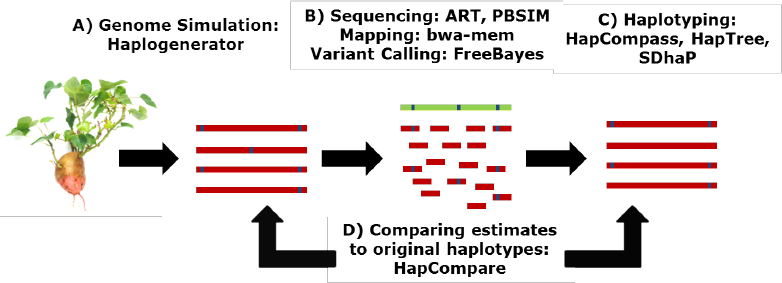
*Haplosim* pipeline to generate, estimate and evaluate haplotypes. Random genomes and haplotypes are produced by *Haplogenerator*, from which NGS reads are simulated and mapped backed to the reference. SNPs are called at the next step and the haplotypes estimated by HapTree, HapCompass and SDhaP. The estimates are compared to the original haplotypes by *hapcompare*.

### 2.1. Polyploid haplotyping software

Currently, three optimization based haplotyping algorithms are available for polyploids: HapCompass [39, 43], HapTree [44] and polyploid SDhaP [40]. We explain each method assuming a genomic region containing *l* heterozygous SNP sites *s*_1_, *s*_2_,…,*s_l_*, a ploidy level equal to *k*>2, and an NGS dataset consisting of *m* (paired-end) reads. We define a “fragment” as the sequence of the determined alleles at the heterozygous sites within a (pairedend) read. For simplicity, we focus on the most prevalent type SNPs, bi-allelic SNPs, for which the alleles can be represented by ’0’ (the reference) and ’1’ (the alternative).

**a) HapCompass:** Aguiar and Istrail (2013) extend their graphical haplotype estimation approach for diploids [39], by constructing the polyploid *Compass graph*, which has *k* nodes for each variant site, *s_i_*, of a *k*-ploid corresponding to the *k* alleles at that site [43]. To each SNP pair, *s_i_*, *s_j_*, that is covered by at least one of the *m* fragments, the phasing with the largest likelihood is assigned by a polyploid likelihood model, conditional on the covering fragments and assuming a fixed base calling error rate. *k* edges are accordingly added to the Compass graph between the nodes at *s_i_* and *s_j_* sites, representing the *k* homologues covering *s_i_* and *s_j_* and weighted by their likelihoods. A global *minimum weighted edge removal (MWER)* algorithm is applied to detect and eliminate edges with conflicting phasing information from the Compass graph with the aid of auxiliary *Chain graphs*. Each Chain graph takes a set of variants that make a cycle in the Compass graph and detects phasing conflicts within the cycle, if present. The MWER algorithm then tries to resolve the conflicts by eliminating a number of edges with minimum total weight corresponding to the least likely phasings. Finally, the most likely haplotypes are found over the full set of SNPs from the conflict free Compass graph by finding *k* disjoint maximum spanning trees through an efficient greedy algorithm, corresponding to the *k* most likely homologues covering the *l* SNP sites.

**b) HapTree:** The HapTree algorithm, developed by Berger et al. (2014) [44], builds a tree representing a subset of likely phasing solutions for *l* heterozygous sites, and tries to find the most probable path from *s*_1_ to *s_l_* within the tree as the best phasing, by calculating the probability of each phasing conditional on the *m* fragments assuming a fixed base calling error rate. Nevertheless, as an exhaustive search over the full tree of all solutions would be computationally prohibitive, the tree is built and extended site by site with branching and pruning to greedily eliminate the (relatively) low probability paths from the final tree. In doing so, HapTree calculates the relative probabilities of the haplotypes at each extension using the relative probabilities of the haplotype at the previous extension that survived the branching and pruning, taking the error model into account.

**c) SDhaP:** The third algorithm, polyploid SDhaP designed by Das and Vikalo (2015), is a semi-definite programming approach that aims to find an approximate MEC solution by a greedy searching of the space of all possible phasings from *s*_1_ to *s_l_* [40]. The algorithm starts with random initial haplotypes, and tries to find the MEC solution by making changes to these initial haplotypes according to a gradient-descent method. To this end, the MEC problem is reformulated as a semi-definite optimization task and preliminary solutions are obtained in polynomial time by exploiting the sparseness of overlaps between fragments for efficient implementation. These preliminary estimates are subsequently refined by greedy flipping of the alleles in the estimated homologues to further reduce the MEC, if possible. By this flipping, SDhaP allows making changes to the dosages of the alternative alleles estimated during variant calling, which could sometimes lower the error correction score. Therefore, the dosages of corresponding SNPs in the SDhaP estimates and original haplotypes could differ, in contrast to the estimates produced by HapTree and HapCompass.

### 2.2. Simulation of polyploid genomes and NGS reads

**a) haplogenerator.** We developed the command-line tool *haplogenerator* for generating artificial genomes and their haplotypes with desired characteristics. Specifying an indexed fasta file as input, one can apply with *haplogenerator* random insertions, deletions and mutations to the input sequence according to a chosen stochastic model to produce modified fasta files for each of the *k*-genomes of a *k*-ploid individual. A separate haplotype file is also made containing the phasing of the generated variants. In the haplotype file, the reference and alternative alleles are numerically coded, assigning 0 to the nucleotide present in the input reference and the following integers to the alternative alleles. The coordinate of each variant on its contig is also specified within the haplotype file.

The random indel and mutation sites are scattered across the input genome according to a selected built-in stochastic model for the distance between consecutive variations, or alternatively by sampling with replacement from a given empirical distribution for this distance. At each position *i*, possible alternative alleles are generated: for indels, an inserted or deleted nucleotide; for mutations, nucleotides other than the reference (or just one nucleotide for obtaining bi-allelic SNPs). Next, a dosage *d_i_* is assigned to the alternative allele based on the ploidy *s_i_*, according to user-specified probabilities for *d_i_*=1 to *d_i_*=*k*, and *d_i_* out of *k* homologues are selected randomly to get their allele at *s_i_* changed to an alternative. To account for linkage disequilibrium, we imposed an additional step after this dosage assignment to reassign the alternative alleles at each *s*_*i*+1_, *i* = 1, 2,…, *n* – 1, to the homologues containing the alternative alleles at *s_i_* (as much as the numbers of alternative alleles, i.e. the dosages, at *s_i_* and *s*_*i*+1_ allow), with a reassignment decision made independently for each site from *s*_2_ to *s_n_* with an arbitrary probability set to 0.4.

**b) Simulation of NGS reads.** We used the technology specific simulator ART [46] to generate paired-end reads from Illumina MiSeq and HiSeq 2500 technologies [51, 52], and PBSIM [47] to simulate Circular Consensus Sequencing (CCS) reads from Pacific BioScience [52, 53]. The average length of single Illumina reads was set to the maximum allowed by ART (125*bp* and 250*bp* for HiSeq 2500 and MiSeq, respectively), and the average read length of CCS was set to 1*kb* for PBSIM, with the read lengths following the built-in distributions derived from empirical data for each technology.

Each homologue was “sequenced” separately, and the reads were combined to simulate the output of real sequencing apparatus. Average sequencing depths were specified for each homologue to obtain the desired average total depth equal to *k* times the per homologue depth, with *k* being the ploidy. Both ART and PBSIM consider a discrete uniform distribution with the user-set mean for the depth at each position, and hence the standard deviation of the total depth was dependent on the average depth per homologue, c, and equal to 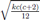) for a *k*-ploid.

The choice of the sequencing strategies in our study was based on the efficiency and performance of the available techniques [52], as well as their practical convenience. In particular, we did not simulate Continuous Long Reads (CLR) of PacBio as they yielded a high false SNP discovery rate at the variant calling stage, which gravely degraded the performance of the haplotyping methods, nor did we use single-ended reads of Illumina as preliminary assessment showed that they produce low quality estimates with a large number of gaps in the solution.

**c) Simulation of polyploid datasets.** In order to simulate realistic polyploid genomes, we chose tetraploid potato (*S. tuberosum*) as the model organism, due to the availability of a reference genome [15] as well as NGS data containing genomic variation of 83 diverse cultivars [45]. The sequence of chromosome 5 from PGSC v4.03 DM draft genome [15] was used as the template sequence for haplogenerator. We selected random contiguous regions with 20*kb* length from this template to be used as references for simulating genomes. In selecting the references, we rejected 20*kb* regions of the template that contained more than 20% undetermined nucleotides, and omitted these undetermined sites (denoted by ’*N*’ in PGSC v4.03 DM sequence) before introducing mutations in the accepted regions. As the length of many genes falls below 20*kb*, choosing references with this size allows us to evaluate haplotype estimation for amplicon sequences covering a complete gene. Random bi-allelic SNPs were introduced in each reference to produce synthetic tetraploid genomes according to the built-in *lognormal* model of haplogenerator, with the mean and the standard deviation of the log-distance between the SNPs being set to 3.0349 and 1.293, respectively, corresponding to an expected SNP frequency of 1 per 21*bp* with a standard deviation of 27*bp*. The distribution of the dosages, *d_i_*, was similarly set equal to that from [45], with percentages equal to 50%, 23%, 14% and 13% of simplex (*d_i_*=1), duplex, (*d_i_*=2), triplex (*d_i_*=3) and quadruplex (*d_i_*=4) SNPs.

In order to investigate the effect of library preparation, we considered various insert-sizes for paired-end Illumina reads, namely end-to-end insert-sizes of 235, 300, 400, 500, 600 and 800*bp* with HiSeq 2500 and 400, 450, 500, 600 and 800*bp* with MiSeq. For evaluation of the effect of sequencing depth on haplotyping, 2×,5×,8×,10×,20×,22×,25×,28×,30× and 35× average coverages were considered per homologue for each of these insert-sizes.

To investigate the effects of genome characteristics, the ploidy level, the dosage of different homologues and the SNP density on the quality of haplotype estimation, additional genomes were generated in a similar way by haplogenerator. Considering the same proportion of simplex and duplex SNPs, i.e. SNPs with dosages equal to 1 and 2, respectively, as in [45] and considering equal proportions for the dosages higher than 2, we simulated genomes with 3*n*, 4*n*, 6*n*, 8*n*, 10*n* and 12*n* ploidy levels to investigate the effect of the ploidy, and simulated modified tetraploid genomes that contained only two distinct homologues with simplex and triplex dosages to investigate the effect of similarity between the homologues on haplotype estimation. While these scenarios assume a SNP-density model valid for *S. tuberosum*, they still can show the pattern by which ploidy level and similarity between the homologues influence the quality of haplotyping.

Finally, tetraploid genomes with SNP densities lower than that of the highly heterozygous *S. tuberosum* [45] were simulated to observe the effect of SNP density, with average frequencies of 1 per 22*bp* to 1 per 110*bp*.

In total, 250 individuals were simulated for each of the above mentioned scenarios by choosing 25 random references from the template and generating 10 genomes with randomly distributed bi-allelic SNPs for each selected reference (Figure 3).

### 2.3. Evaluation of the estimated haplotypes

As several types of error occurring in different steps of the haplotyping pipeline could cause differences between the actual haplotypes and their estimates, we needed several measures of consistency to be able to capture all of them, as summarized in Table 1. These errors include the absence or wrong dosage of original SNPs in the estimates, presence of spurious SNPs, discontinuity of the estimated haplotypes, i.e. presence of gaps between estimated haplotype blocks, and finally wrong extension of homologues leading to incorrect phasing. We also included an extra measure, the *failure rate* (*FR*) for each algorithm, regardless of the quality of haplotype estimation, as it could happen that the haplotyping tools failed to produce any estimate for some of the individuals.

**Table 1.** Summary of the measures used to asses the quality of haplotyping

| Measure | Description | Limitation |
| --- | --- | --- |
| PAR | The average accuracy of phasing for any possible SNP pair | SNPs present in both original haplotypes and estimates, no account of gaps |
| VER* | Number of wrong extensions of homologues with (parts of) other homologues | SNPs present in both original haplotypes and estimates with same dosages, no account of gaps |
| SMR | Proportion of original SNPs missing in the estimates | No account of phasing, false positive SNPs in estimates and dosages |
| IDR | Proportion of SNPs with an incorrect dosage in estimated haplotypes | SNPs present in both original haplotypes and estimates, no account of phasing |
| PPV | Proportion of true SNPs in the estimated SNPs | No account of phasing, missing SNPs, and wrong dosages |
| NGPS* | Number of gaps (interruptions) introduced in the estimated haplotype | No account of phasing, missing, wrong dosages and false positives |
| FR | Rate of failure for an algorithm | No account of the quality of the estimates |

\*: measure normalized by the size of original haplotypes, i.e. the number of original SNPs

Some of the SNPs originally simulated were absent in the output set of each simulation, either due to mapping and variant calling errors or due to their not being phased by the algorithms, and therefore could not be included in the comparisons of true and estimated phasings. Instead, their proportion was calculated as *SNP missing rate* (*SMR*) and considered as the first measure of estimation quality. Similarly, spurious SNPs in the estimates were not included in the comparisons and the *Positive Predictive Value* (*PPV*) or the precision of the estimated SNPs was calculated as the proportion of genuine SNPs among the estimated SNPs as the next measure of estimation accuracy. The *incorrect dosage rate* (*IDR*) was also calculated as the proportion of SNPs that had an incorrect dosage in the estimated haplotypes in the set of SNPs that were common between the original haplotypes and the estimates.

Having excluded the missing and spurious SNPs, the *Pairwise Phasing Accuracy Rate* (*PAR*) [54] was computed as the proportion of all heterozygous SNP pairs for which the estimated phasing was correct. This measure captures the errors caused by chimeric elongation of the homologues during haplotype estimation, i.e. the elongation of a homologue by (part of) another homologue, as well as errors caused by incorrect dosage estimation.

One way to calculate the accuracy of phasing for more than just two SNPs is to consider the phasing accuracy rate for groups of three SNPs, four SNPs, etc. However, the phasing accuracies will no longer be independent for the groups of SNPs that have more than one SNP in common, leading to biased estimates of the accuracy rates. Instead, we calculated the *Vector Error Rate* (*VER*), also called the switch error rate, defined as the number of times a homologue is erroneously extended by part of another homologue [44]. Such erroneous extensions are also called switches between homologues, and the measure is equal to two times the number of wrong phasings for pairs of consecutive SNPs for diploids. For polyploids, the measure is calculated by finding the minimum number of crossing-overs needed to reconstruct the true haplotypes from the estimates [44]. To be able to compare of VER for different ploidy levels, genome lengths and SNP densities, we normalized it by the number of originally simulated SNPs as well as the ploidy level for each individual. The SNPs with a wrong estimated dosage of the alternative allele were omitted before applying this measure, as otherwise the true haplotypes could not be reconstructed by simple switching of the estimated homologues without considering allele flips from 0 to 1 or *vice versa*.

The last measure of estimation quality that we used was the *number of gaps per SNP* (*NGPS*) in the estimates, as the simulated continuous haplotypes can be broken into several disconnected blocks, causing gaps in the estimated haplotypes. This phenomenon happens if the connection between SNPs is lost due to low sequencing coverage or sequencing/variant calling errors at certain sites. Therefore, we calculated the number of break points, i.e. gaps, in the estimates (equal to the number of disjoint blocks minus one), and normalized it by the total number of simulated SNPs for each individual for the same reasons as for VER. In case gaps were present in the estimates, we calculated the other measures separately for each estimated block and reported the weighted average of the block-specific measures, weighted by the number of compared SNPs in each estimated block, or the number of possible pairwise phases in case of PAR.

Finally, the computational complexity of each of the haplotyping algorithms was considered as a function of sequencing coverage, insert-size, ploidy, SNP density and homologue dosages. The applied haplotyping methods are memory intensive methods, increasingly consuming system resources with time, sometimes up to tens of gigabytes of virtual memory. To run the methods on a system with shared resources, and considering the fact that the algorithms require an increasing amount of virtual memory with time, a time limit of 900 seconds (2000 seconds for the analysis with various levels of ploidy) was imposed on each haplotyping algorithm, after which the algorithms were externally halted and the estimation considered a failure. This amount of time was deemed reasonable considering the 20*kb* length of the simulated genomic regions, and the number of time-out events was added to the number of times each algorithm failed to estimate any haplotypes due to the occurrence of internal errors. Total FRs are thus also reported for each haplotyping scenario.

### 2.4. Comparison of haplotyping algorithms

In order to compare the overall performance of the three haplotype estimation methods, we built three linear regression models with the mentioned quality measures as response and the haplotyping method as predictor, considering sequencing depth, sequencing technology and the paired-end library size as covariates in the model. As each of the simulated genomes was haplotyped simultaneously by the three estimation methods, the effect of the genome on the estimation quality was incorporated as a random effect in the model. Similarly, as 10 genomes were generated from each of the 25 randomly selected references, the effect of the common reference was added as the second random component to the model.

For each quality measure, a complete-case analysis was performed, including only the results of those simulations for which all the three estimation methods reported some value. The models were estimated by Restricted Maximum Likelihood (REML) [55] using the *lmer* function from the package *lme4* [56] in R 3.2.2 [57].

## 3. Results and Discussion

### 3.1. Haplogenerator produces realistic genomes

In order to investigate the compatibility of the simulated 20*kb S. tuberosum* genomic regions with the real regions sequenced by uitdewilligen et al.(2013) [45] in terms of the density of bi-allelic SNPs, we obtained quantile plots (QQ-plots) of the distances between consecutive SNPs *s_i_*, *s*_*i*+1_, generated by the applied lognormal model versus the distances between consecutive bi-allelic SNPs (within the same RNA-capture region) in the combined data of 83 diverse cultivars from [45]. As shown in Figure 2, the two empirical distributions match well enough, although the distribution of real SNPs seems to have a heavier tail than lognormal (accounting for less than 2% of the total number of real bi-allelic SNPs). This heavier tail is plausibly explained by the presence of highly conserved regions in real genomes, subject to natural and artificial selection pressure, and the limitations of the RNA-capture sequencing used in [45] to detect all of the SNPs within a captured region with the applied sequencing depth.

**Figure 2.**
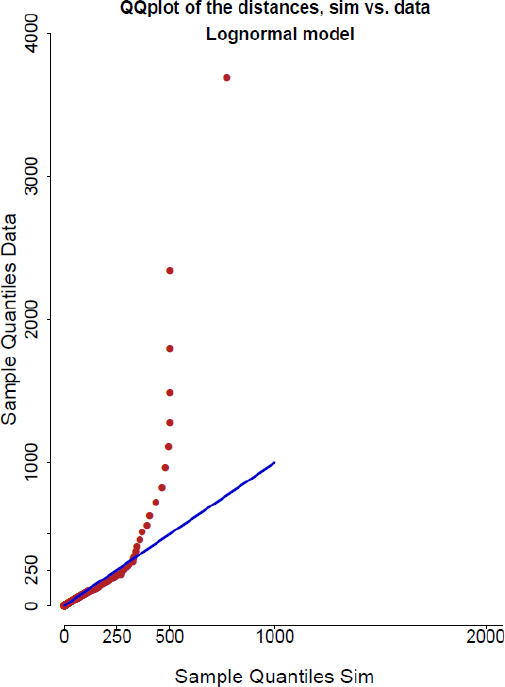
Quantile plots for the distances between successive SNPs obtained from simulation by *haplogenerator* using the lognormal distance model (horizontal axis) versus the one obtained from the data of 83 potato cultivars of Uitdewilligen et al. (2013) [45]. The two distributions match well, though a heavier tail is observed for the data of Uitdewilligen et al. (2013), accounting for less than 2% of the SNPs.

**Figure 3.**
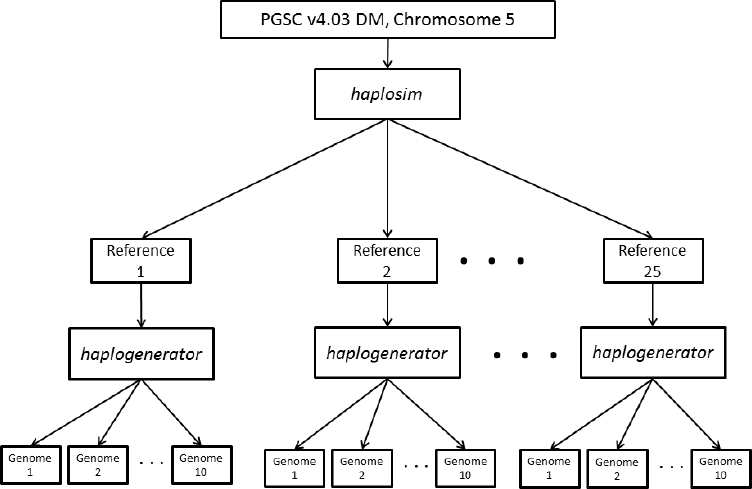
The schematic diagram of simulation for each considered scenario. 10 references of length 20kb are chosen from the draft sequence chromosome 5 [15], from each 10 polyploid genomes are simulated containing bi-allelic SNPs randomly distributed according to the lognormal distance model, to obtain datasets of size 250.

The proportions of simplex to quadruplex SNPs, i.e. the dosage proportions, were also almost identical as those obtained from Uitdewilligen et al. (2013) [45] (Section 2.2-c).

### 3.2. Sequencing depth is the major determinant of haplotyping quality

An important goal of the simulations was to observe to what extent sequencing strategies influence haplotyping results, because of the practical importance in setting up experiments and choosing a technology. As different technologies rely on different library preparation and nucleotide calling methods, their output is often different in terms of the average read length and sequencing error profile. Besides, the sequencing depth, average read length and paired-end insert-size can vary according to the user’s requirements with the same technology. We found that the performance of all three haplotyping methods was considerably affected by the sequencing strategy, most notably by the sequencing depth.

Regardless of the used sequencing technology and the insert-size, a sequencing depth between 5-20× per homologue is required to obtain results satisfactory in terms of haplotype accuracy (PAR) and completeness (SMR) (Figure 4-a,b). Both improve continuously with sequencing depth, but flatten out at 15×. A notable exception is HapTree, of which the increased failure rate (FR) at higher depths is reflected in a worse completeness (increasing SMR). Other quality measures (VER, PPV, IDR, NGPS) were not substantially influenced by sequencing depth at depths higher than 5× per homologue.

**Figure 4.**
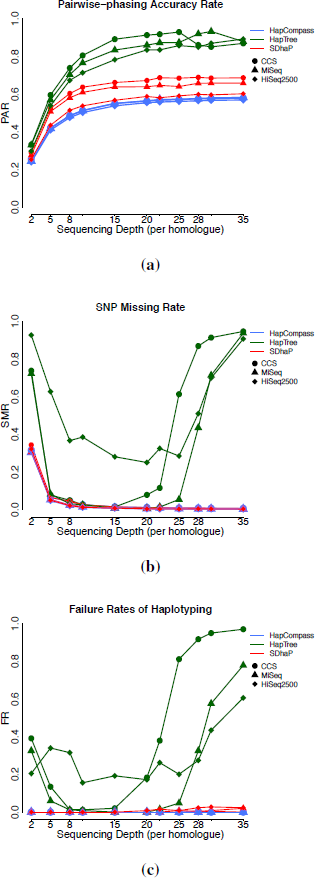
Plots of haplotype estimation quality measures: (a) SMR, (b) PAR and (c) FR against sequencing depth per homologue using HapCompass (blue), HapTree (green) and SDhaP (red), for simulated 20*kb* tetraploid *S. tuberosum* genomes. Sequencing was performed in silico for paired-end MiSeq (triangle) and HiSeq 2500 (rhombus) with 800*bp* insert-size, as well as for PacBio-CCS of 1*kb* length (circle).

HapTree’s failure rate (FR, Figure 4 c) was rather high for low and high sequencing depths. At lower depths, less than 2× per homologue, there is not enough information available for effective branching and pruning of the solution tree and time-out errors result in failures. In contrast, for high sequencing depths the relative likelihood values often become very small, due to the presence of many terms in the likelihoods of partial haplotypes, making a meaningful comparison impossible (a penalty for not using normalized Bayesian posteriors). This problem will be discussed further in Section 3.8.

As sequencing depth is an important factor in determining the total cost of sequencing, these results show that extra cost can be avoided by choosing a moderate sequencing depth without sacrificing considerable haplotyping accuracy.

### 3.3. Large insert paired-end reads are competitive with long reads

In addition to sequencing depth, the insert-size of paired-end reads and the employed sequencing technology can have an impact on the estimation quality. These are also important factors to specify when designing a sequencing experiment, as they influence cost, throughput and quality. To quantify their effects, we simulated NGS reads for HiSeq 2500, MiSeq and CCS technologies at each sequencing depth, and simulated paired-end reads with various insert sizes for HiSeq 2500 and MiSeq (Section 2.2-b).

Our results show that at the same sequencing depth, increasing the insert-size of paired-end reads was not markedly influential on the overall quality of haplotyping (Supp. Figure 1), except for the number of gaps that was expectedly reduced with larger inserts. Moreover, similar estimation qualities were obtained using the long (1*kb*) PacBio CCS reads and paired-end Illumina reads with a large insert-size (800*bp*) (Figure 4). At the same depth, the paired-end reads contain basically the same phasing information as the long reads.

Although libraries with large inserts are costly and difficult to obtain, they may be still easier to generate than long continuous reads and therefore can be a competitive option for designing haplotyping experiments.

### 3.4. HapTree is the most accurate method, but often fails

Different haplotyping algorithms yield different estimates for the same individual. With simulated individuals, it is possible to compare the quality of these estimates as the haplotypes are known *a priori*. To this end, we used linear regression models relating the performance measures to the algorithms used (Section 2.4).

Table 2 shows the 99% confidence intervals for the effects of estimation method and sequencing technology on the haplotyping accuracy for the tetraploid genomes, with HapCompass on PacBio data taken as the baseline. HapTree is significantly more accurate (higher PAR and lower VER) than the other methods, but less complete (higher SMR) due to its frequent failure (Figure 4-c). SDhaP yields slightly, but significantly, worse dosage estimates (higher IDR). There was no significant relation between the method used and the continuity of haplotype estimates (NGPS) or precision of the SNPs (PPV). Finally, Illumina reads perform only very slightly worse than PacBio, although in a couple of cases still significant at *α*=0.01.

**Table 2.**
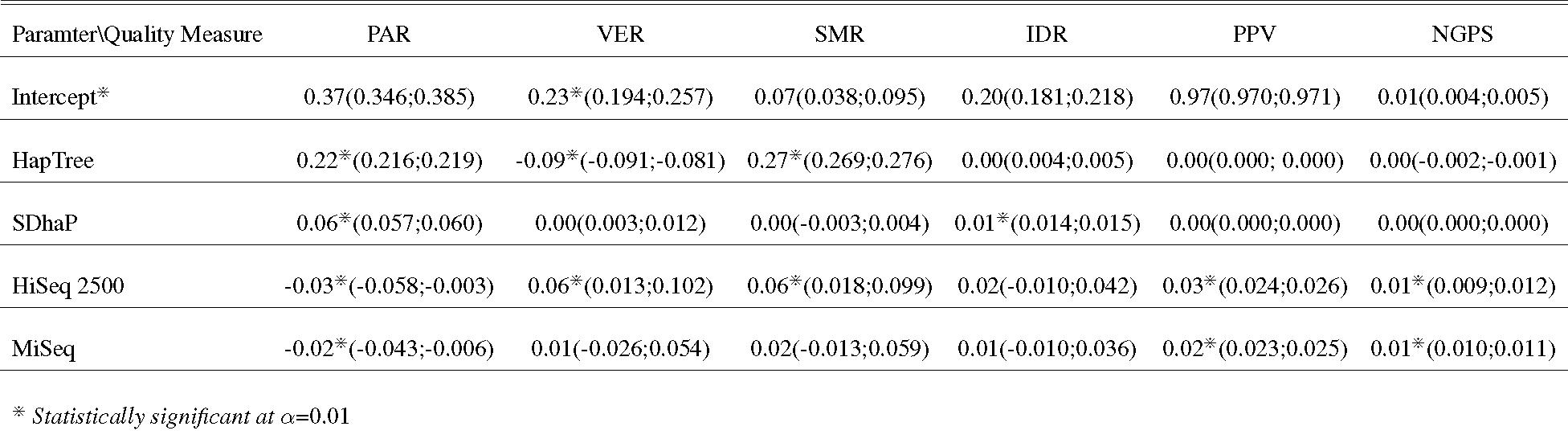
Point estimates and 99% confidence intervals for the effects of haplotyping and sequencing methods on the haplotyping quality measures

Overall, these results confirm that HapTree is the most accurate method when it does not fail, and that Illumina and PacBio reads offer very similar performance.

### 3.5. Similarity between homologues eases haplotyping with Illumina

Similarity between homologues can have a large effect on haplotyping. This similarity often occurs when random mating is violated, e.g. in inbred or isolated populations. To investigate this, we simulated simplex-triplex individuals, i.e. tetraploid individuals consisting of two different genomes with dosages of 1 and 3. We generated paired-end MiSeq and HiSeq 2500 reads (800*bp* insert-size), as well as 1*kb* CCS read of PacBio, and evaluated the estimated haplotypes.

On this data, the performance of HapCompass and HapTree with Illumina reads improves over the original simulation, while the performance of SDhaP deteriorates significantly (Figure 5). In particular, the similarity between homologues resulted in a decreased accuracy for SDhaP (Figure 5-a, PAR around 0.2) and incorrect dosage estimates for more than half of the SNPs (Figure 5-c, IDR of 0.55), regardless of the sequencing method.

**Figure 5.**
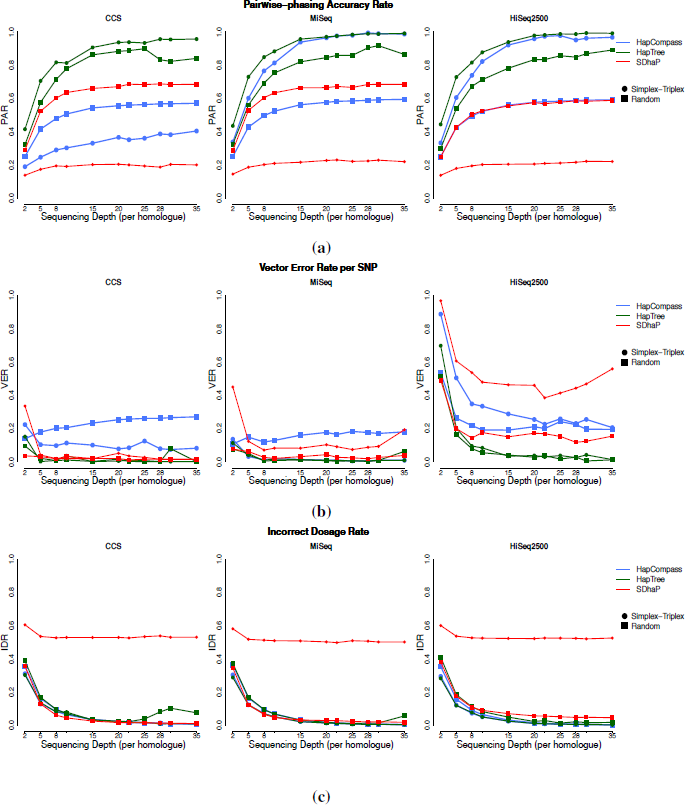
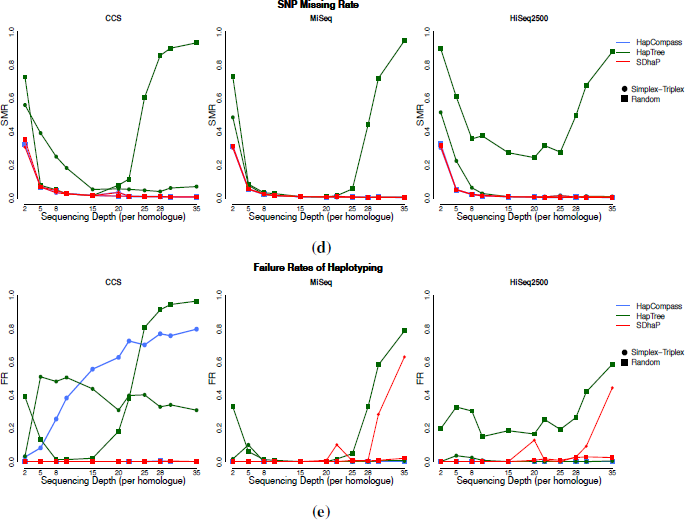
Plots of haplotype estimation quality measures: (a) PAR, (b) VER, (c) IDR, (d) SMR and (e) FR against ploidy level using HapCompass (blue), HapTree (green) and SDhaP (red), for simulated 20*kb* simplex-triplex tetraploid genomes (circle) compared to genomes with random haplotype dosages (square). Sequencing was performed in silico for paired-end HiSeq 2500 reads with 800*bp* insert-size.

These results demonstrate the differences between the MEC (Minimum Error Correction) approach to haplotyping and other approaches. MEC is sensitive to (local) similarities between homologues, as they lead to approximately identical MEC scores for several different phasings and cause SDhaP to report a suboptimal solution. In contrast, the performances of HapCompass (MWER approach) and HapTree (relative likelihood approach) improve, at least when using Illumina sequencing (Figure 5-a,b). Having more similar fragments simplifies construction of the maximum spanning tree in the Compass graph and makes the branching and pruning of the solution tree of HapTree more accurate by enhancing the relative likelihoods of correct partial phasings. No improvement was observed, however, with CCS reads, due to increasing failure rates (FR, Figure 5-e) caused by time-out errors.

These results show that the underlying algorithms lead to different sensitivities to homologue similarity, with MEC-based approaches yielding incorrect results and other methods demanding increasing computation time.

### 3.6. SNP density is mostly influential on the continuity of haplotype estimates, but not on the other measures of estimation quality

In genomes with a lower SNP density than the highly heterozygous potato, *S. tuberosum*, overlaps will occur less often between the fragments, which can influence the quality of haplotyping. To determine the effect of SNP density, we simulated tetraploid genomes with average SNP densities ranging from 1 SNP per 22 base pairs, the average density for potato, to 1 SNP per 110 base pair, and estimated the haplotypes using Illumina paired-end reads with an insert-size of 800*bp*, as well as 1*kb* CCS reads of PacBio, at a sequencing depth of 15× per homologue. Increasing numbers of gaps (NGPS, Figure 6-a) and a decrease in completeness (SMR, Figure 6-b) were observed in the estimated haplotypes at lower densities for all three haplotyping methods. The effect of the SNP density was, however, not manifest on the other haplotyping quality measures (Supp. Figure 5). In summary then, SNP density mostly influences the continuity of haplotype estimates.

**Figure 6.**
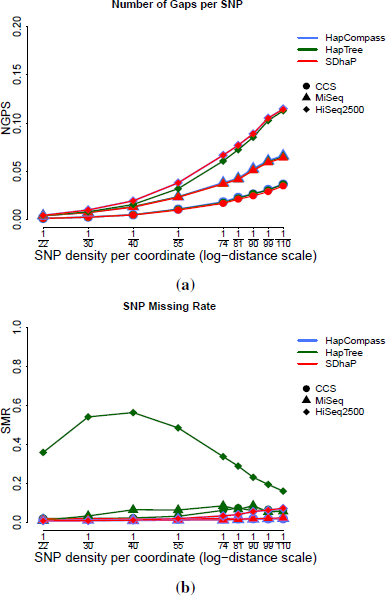
Plots of haplotype estimation quality measures: (a) NGPS, (b) SMR against SNP density (at logarithmic distance scale) using HapCompass (blue), HapTree (green) and SDhaP (red), for simulated 20*kb* tetraploid *S. tuberosum* genomes. Sequencing was performed in silico for paired-end MiSeq (triangle) and HiSeq 2500 (rhombus) with 800*bp* insert-size, as well as for PacBio-CCS of 1*kb* length (circle), at a depth of 15×.

### 3.7. At higher ploidy levels, HapCompass is the best method to use

In order to investigate whether our findings for tetraploid genomes hold for other ploidy levels, we performed simulations with ploidy levels of 3-12 (Section 2.2-c). We simulated paired-end HiSeq 2500 reads with an insert-size of 800*bp*, as it gave high quality estimates in tetraploids and was more practical than the competitive sequencing options, at 5×, 15× and 20× sequencing depths per homologue.

The accuracy of HapTree and SDhaP decreases markedly with increasing ploidy level, up to 30% for 12*n* (PAR, Figure 7-a), while the performance of HapCompass remained stable. Likewise, the completeness of HapTree decreased (SMR, Figure 7-b) and failure rates for both HapTree and SDhaP increased (Figure 7-c). Although the performance of the methods at each ploidy level was relatively better at higher sequencing depths, the deterioration of the haplotype estimation quality followed a similar pattern with the increase in ploidy, regardless of the depth.

**Figure 7.**
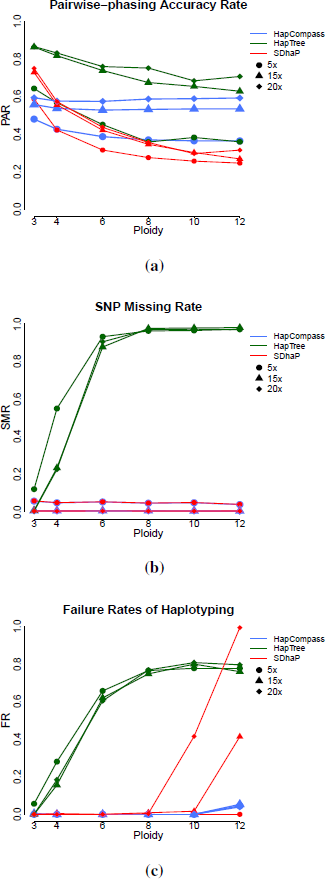
Plots of haplotype estimation quality measures: (a) PAR, (b) SMR and (c) FR against ploidy level using HapCompass (blue), HapTree (green) and SDhaP (red), for simulated 20*kb* 3*n*, 4*n*, 6*n*, 8*n*, 10*n* and 12*n* genomes. Sequencing was performed in silico for paired-end HiSeq 2500 with 600*bp* insert-size. Three sequencing depth were used per homologue: 5x (circle), 15x (triangle) and 20x (rhombus).

Overall, none of the haplotyping methods is equipped to deal with high levels of ploidy: either they break down (HapTree, SDhaP) or are inaccurate (HapCompass).

### 3.8. HapTree has in general the highest memory consumption and computation time compared to the other methods. Both factors depend on the length of the genome, ploidy level and sequencing depth for HapTree

Computational efficiency is an important feature of every complex algorithm, such as the haplotyping algorithms discussed in this paper. Therefore, we measured the memory and time consumption of each algorithm for various ploidy levels, sequencing coverages and genome lengths. Using HiSeq 2500 and paired-end libraries with an insert-size of 800*bp* for the simulation of sequencing, we tested the effect of sequencing depth with tetraploid individuals and genomes of length 10*kb*, and the effect of genome length with tetraploid individuals sequenced at an average depth of 10× per homologue. Other settings were the same as for *S. tuberosum* (Section 2.2), for each condition generating 50 individuals from 50 randomly selected regions, i.e. one individual per region, with a time limit of 7200 seconds. Fixing the depth to 10× per homologue and the genome length to 10*kb*, the effect of ploidy was also investigated in a similar manner.

The analyses were run on multicore 2.6 GHz Intel-Xeon processors. For each the total CPU-time and physical memory consumption was measured using the Unix getrusage routine. HapTree clearly consumed most time and memory resources (Figure 8), increasing with genome length (Figure 8-a,b) and ploidy level (Figure 8-c,d). This increase was much less for HapCompass and SDhaP.

**Figure 8.**
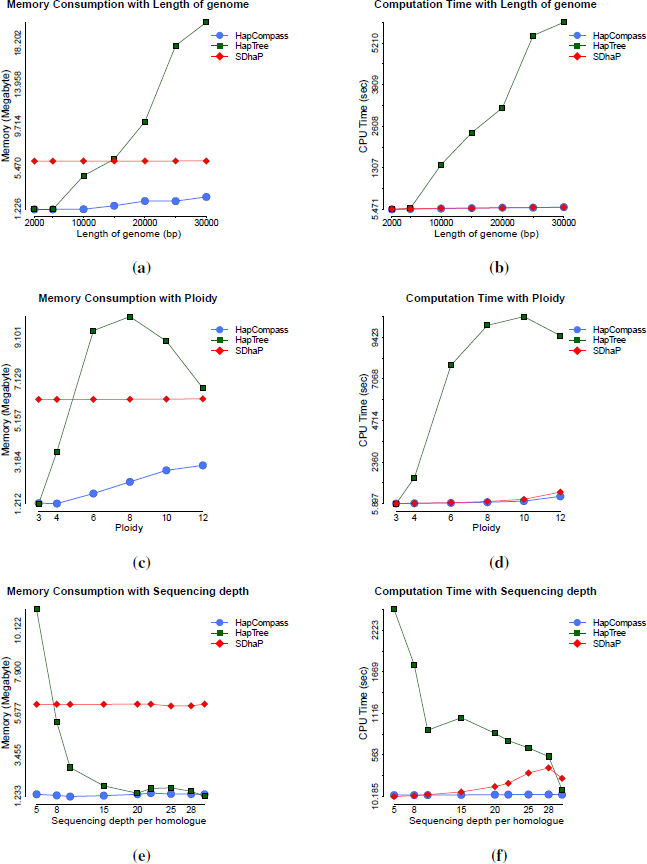
Plots of physical memory consumption (in Megabyte) with: length of genome (in base-pair) (a), ploidy (c) and sequencing depth (e), and plots of computation time (in seconds) with length of genome (in base-pair) (b), ploidy (d) and sequencing depth per homologue (f), for three haplotype estimation softwares: HapCompass (blue-circle), HapTree (green-square) and SDhaP (red-rhombus).

While sequencing depth was less influential, Haptree used much more time and memory at the lowest sequencing depth, 5× per homologue, falling rapidly with an increase in depth up to 10× per homologue and remaining almost constant afterwards up to a depth of 20× per homologue (Figure 8-e,f). At the depths of higher than 20× per homologue, the computation time fell rapidly again due to the premature failure of the algorithm as discussed in Section 3.2.

## 4. Conclusion

We evaluated three algorithms for single individual haplotype estimation in polyploids: HapCompass, HapTree and SDhaP, and investigated the effects of sequencing technology, insert size and sequencing depth on the estimation quality using several measures of quality (Table 1) and through extensive simulation experiments.

Our results show that HapTree can produce the best triploid and tetraploid haplotype estimates, followed by SDhaP and HapCompass. HapCompass is the best method to use with ploidy levels ≥6n, although its performance is not very good in the absolute sense. We showed that sequencing depth was the most important factor determining the quality of haplotype estimation, and paired-end short reads of Illumina with a large insert can perform as well as long CCS reads of the same total size now possible with PacBio. For accurate haplotyping, we therefore suggest an average depth of between 5-20× per homologue with paired-end reads and an insert-size of 600-800*bp*. In addition to the estimation quality, we investigated computation time and memory consumption of each algorithm under various settings to compare their efficiency. We showed that on average, HapTree requires the most computation time and memory, and its use of resources is highly dependent on the length of the genomic region, the ploidy level and the sequencing depth. Combined with the frequent failure to complete the estimation, this raises difficulties for applying HapTree on practical problems where the aim is to reconstruct long-range haplotypes.

Our findings show that while state-of-the-art haplotype estimation algorithms produce promising results for triploid and tetraploid organisms over a limited genomic region, their performance rapidly decreases at higher ploidy levels and their resource use prohibits application to large genomic regions. The probability-based algorithm of HapTree produces the most accurate estimates but also requires the most computation time and memory. We believe it is worth investigating whether HapTree can be made robust when faced with larger problems while maintaining its accuracy, e.g. using a divide-and-conquer approach or by adjusting the branching and pruning parameters according to the length of the genome, the ploidy level and the sequencing coverage. The variant calling error model could also be upgraded to be specific to the applied sequencing strategy and technology. Finally, the performance of haplotyping methods on individual organisms could be greatly improved if it could also incorporate parental and sib information if available, e.g. mapping populations relevant to plant and animal breeding studies. Such algorithmic improvements will prove essential to help understand the complex genetics found in many polyploid organisms and, in the long run, to better understand the rules governing genome organization.

## Key points

- Haplotyping accuracy for polyploids depends mostly on sequencing depth, with 5-20× giving optimal results.
- Paired-end Illumina reads with large insert-sizes are competitive with PacBio CCS reads at reduced cost as long as the quality of haplotype estimation is concerned.
- For ploidy levels up to six, HapTree is the most accurate method and can produce high quality estimates; above that, HapCompass is the best method to use although its estimation quality is not so high in the absolute sense.
- In spite of its high accuracy, HapTree breaks down for long genomic regions and high ploidy levels, often failing to converge within a reasonable amount of time.
- Improvements are needed to allow accurate haplotyping in polyploids in large genomic regions and may be found in adapting parameters to the problem at hand and in using sibship and parental information if available.

## Acknowledgments

This work was funded by the graduate school Experimental Plant Sciences (EPS) of the Wageningen University and Research Centre. The authors wish to thank Herman J van Eck for his input and valuable discussions regarding the data from Uitdewilligen et al. [45].

## Author Contributions

RF, CM, EM and DdR designed the study, revised and approved the manuscript. EM developed the simulation pipeline and performed the analyses.

---

**Supp Figure 1. SuppFig1_InsertSizes.pdf:** Plots of all of the haplotype accuracy measures against sequencing depth, using paired-end MiSeq and HiSeq 2500 reads with various insert-sizes.

**Supp Figure 2. SuppFig2_Figure4_All_Quality_Measures.pdf:** Plots of all of the haplotype accuracy measures against sequencing depth for tetraploid genomes, using paired-end MiSeq and HiSeq 2500 sequence reads with an insert-size of 800*bp*, as well as PacBio CCS reads of 1kb length.

**Supp Figure 3. SuppFig3_Figure5_All_Quality_Measures.pdf:** Plots of all of the haplotype accuracy measures against sequencing depth, using paired-end MiSeq and HiSeq 2500 sequence reads with an insert-size of 800*bp*, as well as 1*kb* CCS reads of PacBio, comparing random and Simplex-Tripelx haplotype dosages.

**Supp Figure 4. SuppFig4_Figure6_All_Quality_Measures.pdf:** Plots of all of the haplotype accuracy measures against SNP density for tetraploid genomes, using paired-end MiSeq and HiSeq 2500 sequence reads with an insert-size of 800*bp*, as well as PacBio CCS reads of length 1*kb*, at a sequencing depth of 15× per homologue.

**Supp Figure 5. SuppFig5_Figure7_All_Quality_Measures.pdf:** Plots of all of the haplotype accuracy measures against the ploidy level, using paired-end HiSeq 2500 reads with an insert-size of 800*bp*, at sequencing depths of 5×, 15× and 20× per homologue.

